# Computer–Assisted Systematic Chemical–Space Mapping of a First–in–Class Peripherally Restricted α_2A_AR Agonist through Scaffold–Seeded Enumeration

**DOI:** 10.64898/2026.09.16.751790

**Authors:** Cedric Moore, Larry Zhu, Zack Cheng

## Abstract

CC10137 is a first-in-class peripherally restricted α_2A_-adrenergic receptor (α_2A_AR) agonist with broad-spectrum analgesic efficacy and a favorable safety profile. Systematic exploration of the chemical space surrounding first-in-class leads is important for defining series boundaries and guiding continued optimization, but conventional analogue-by-analogue medicinal chemistry samples only a small fraction of the accessible structural space. Here, we used a scaffold-seeded enumeration strategy to expand the chemical space surrounding CC10137 from four SAR-informed seed compounds comprising CC10137 and three closely related structural variants. Application of predefined medicinal chemistry transformation rules in StarDrop generated a virtual library of 16,601,163 unique structures. Morgan fingerprint-based principal component analysis indicated that the library occupied a highly multidimensional structural space involving variation in scaffold substitution, peripheral functional groups, and side-chain composition. A retrospective comparison set of 43 compounds independently designed and experimentally characterized in the earlier CC10137 program represented only approximately 0.00026% of the 16.6-million-member library, yet all 43 were recovered as exact structural matches. Three compounds selected directly from the virtual library retained α_2A_AR binding affinity and agonist potency below 25 nM. Five representative compounds further showed significant anti-allodynic effects in the *in vivo* spared nerve injury model, with inhibition rates ranging from 39.3% to 55.7%. These findings support scaffold-seeded computational enumeration as a practical strategy for systematic chemical-space mapping around a first-in-class lead and for identifying additional pharmacologically active structural regions for further optimization.

## Introduction

α_2A_ adrenergic receptor (α_2A_AR) agonists have established analgesic activity, but their clinical use is limited by central nervous system-mediated adverse effects, including sedation, hypotension, and hypothermia.(*1–6*) However, it had remained unclear whether restricting drug distribution to peripheral tissues could preserve analgesic efficacy while reducing these dose-limiting effects. The development of CC10137 and other peripheral analgesics provided evidence that this approach is feasible.(*7–12*) CC10137 was developed as an orally bioavailable, peripherally restricted α_2A_AR agonist. It is the first peripherally restricted α_2A_AR agonist with broad–spectrum analgesic efficacy and a favorable safety profile, a first–in–class molecule that establishes a new chemical framework for peripherally restricted analgesics.(*8*)

The initial discovery program and the directly related patent disclosed hundreds of analogues within this series.(*13, 14*) However, these compounds represent only a small fraction of the potentially accessible chemical space. For first–in–class molecules, systematic exploration of the surrounding chemical space is critical to the development of second-generation compounds with the potential to achieve greater specificity, improved efficacy, and/or reduced toxicity. Unlike follow–on programs, where established structure–activity relationships provide a navigable starting point(*15*), first–in–class discoveries begin with limited prior knowledge(*16*). The structural landscape is largely uncharted, and the relationship between structural variation and pharmacological properties must be established *de novo*. Every unexplored analogue or substitution pattern represents a potential gap in understanding the structural boundaries and structure–activity relationships of the series.(*17*)

Traditional medicinal chemistry, however, is inherently labor–intensive and time–consuming.(*18*) Even a highly productive team of medicinal chemists may spend years designing, synthesizing, and testing a few hundred analogues.(*19*) A more comprehensive exploration of the surrounding chemical space is therefore difficult to achieve through conventional analogue-by-analogue medicinal chemistry alone. The gap between what is synthetically possible and what can be practically explored within a reasonable timeframe by manual effort has therefore remained a fundamental bottleneck, particularly for first–in–class discovery programs where the need for systematic mapping is greatest. (*20–22*)

Computational approaches provide a complementary means of addressing this limitation by enabling rapid exploration of virtual chemical spaces far beyond what can be synthesized and tested experimentally. Computer-aided molecular design encompasses a range of strategies, including structure-based virtual screening, de novo or generative molecular design, and rule-based chemical-space enumeration. Among these approaches, medicinal-chemistry transformation-based methods are particularly well suited to lead-series expansion because they generate new structures by systematically applying structural modifications derived from established medicinal chemistry precedent, while allowing defined regions of a parent scaffold to be preserved. Such approaches can therefore combine the breadth of computational enumeration with medicinal-chemistry knowledge to explore structurally relevant analogues at a scale that is unattainable through manual design alone.(*8, 23*)

Related transformation-based approaches have been used to construct large virtual chemical spaces and to expand the structural neighborhoods surrounding seed molecules, including DrugSpaceX and ChemBang(*24, 25*). Medicinal chemistry transformation sets derived from literature precedent have been shown to encompass diverse modification classes, including functional-group addition and exchange, linker modification, and ring addition, modification, or removal, and to generate a broad range of chemically acceptable drug-like structures.(*26*) Modern implementations substantially extend the number of available precedented transformations; for example, the StarDrop transformation framework provides approximately 30,000 such rules. Together, these transformations provide broad coverage of medicinal-chemistry-relevant structural modifications.

Following the publication of our study describing CC10137 and its associated medicinal chemistry program, the objective of the present study was to determine whether a scaffold-seeded, rule-based computational enumeration strategy could systematically expand and map the chemical space surrounding this first-in-class scaffold more efficiently than conventional analogue-by-analogue design. Based on the molecular architecture and structure–activity relationships established in the earlier CC10137 program, CC10137 and three closely related structural variants were selected and defined as four SAR-informed seed compounds. The three additional seeds encoded alternative, SAR-supported starting configurations, including variations in linker length and polar-group orientation, as well as linker hopping, thereby reducing dependence on a single starting topology before computational expansion. StarDrop was selected because its transformation-based framework allows a large repertoire of precedented medicinal chemistry modifications to be systematically applied to predefined modifiable regions while preserving structural features considered important for activity. (*26*) The resulting virtual library was subsequently benchmarked against a defined set of 43 compounds comprising the complete Series B and D and Series E-d, which included all dexmedetomidine-related compounds from Series E. These compounds had been independently designed, synthesized, and experimentally characterized during the earlier medicinal chemistry program, providing a direct test of whether the computationally generated space could recapitulate pharmacologically relevant regions previously reached through manual medicinal chemistry. In addition, representative compounds were selected from the generated library for experimental evaluation, including chemical synthesis, α_2A_AR binding and agonist assays, and in vivo testing in the spared nerve injury model. This combined structural and experimental validation strategy allowed us to assess not only whether the computational workflow could recover previously validated medicinal chemistry outcomes, but also whether structures sampled from the expanded chemical space could retain relevant *in vitro* pharmacological activity and *in vivo* anti-allodynic efficacy.

## Results

### Generation of the CC10137-derived chemical space

The chemical-space expansion strategy was guided by the molecular architecture and structure-activity relationships established for CC10137, shown in Figure 1. CC10137 was first decomposed into a fixed receptor-interacting core and several modifiable regions, including the linker, polar, and other variable regions. Because transformation-based molecular generation is dependent on the structural context of the starting molecule, use of CC10137 alone could restrict enumeration to transformations applicable to a single initial topology and substitution pattern. We therefore used three additional, closely related seed structures to provide multiple SAR-informed starting configurations while retaining the conserved α_2A_AR-interacting framework. To capture representative structural features from the major previously explored series, one compound was selected from each of Series B, D, and E based on the medicinal chemistry and SAR knowledge established in the earlier program. These selections were intended to encode complementary structural variations, including changes in linker length, inversion of polar-group orientation, and linker hopping. D12-seed-1 to D12-seed-3 were selected based on SAR from the earlier medicinal chemistry program, including variations in linker length and polar-group orientation, as well as linker hopping. Specifically, Seed-1 and Seed-3 represented alternative linker configurations connecting the dexmedetomidine-related region to the polar moiety, whereas Seed-2 represented inversion of the polar-group orientation. The purpose of the additional seeds was therefore not simply to increase library size, but to reduce starting-structure bias and enable the transformation rules to explore neighboring chemical space from multiple pharmacologically relevant structural configurations. D12-seed-1, D12-seed-2, and D12-seed-3 corresponded to compounds E2, D11, and B3, respectively, which had been previously reported in the CC10137 discovery study(*8*) and therefore represented experimentally established structural variants within the series. Thus, the multi-seed design intentionally combined experimentally established SAR variants providing multiple pharmacologically relevant starting configurations for chemical-space expansion rather than relying on a single molecular topology.

**Figure 1.**
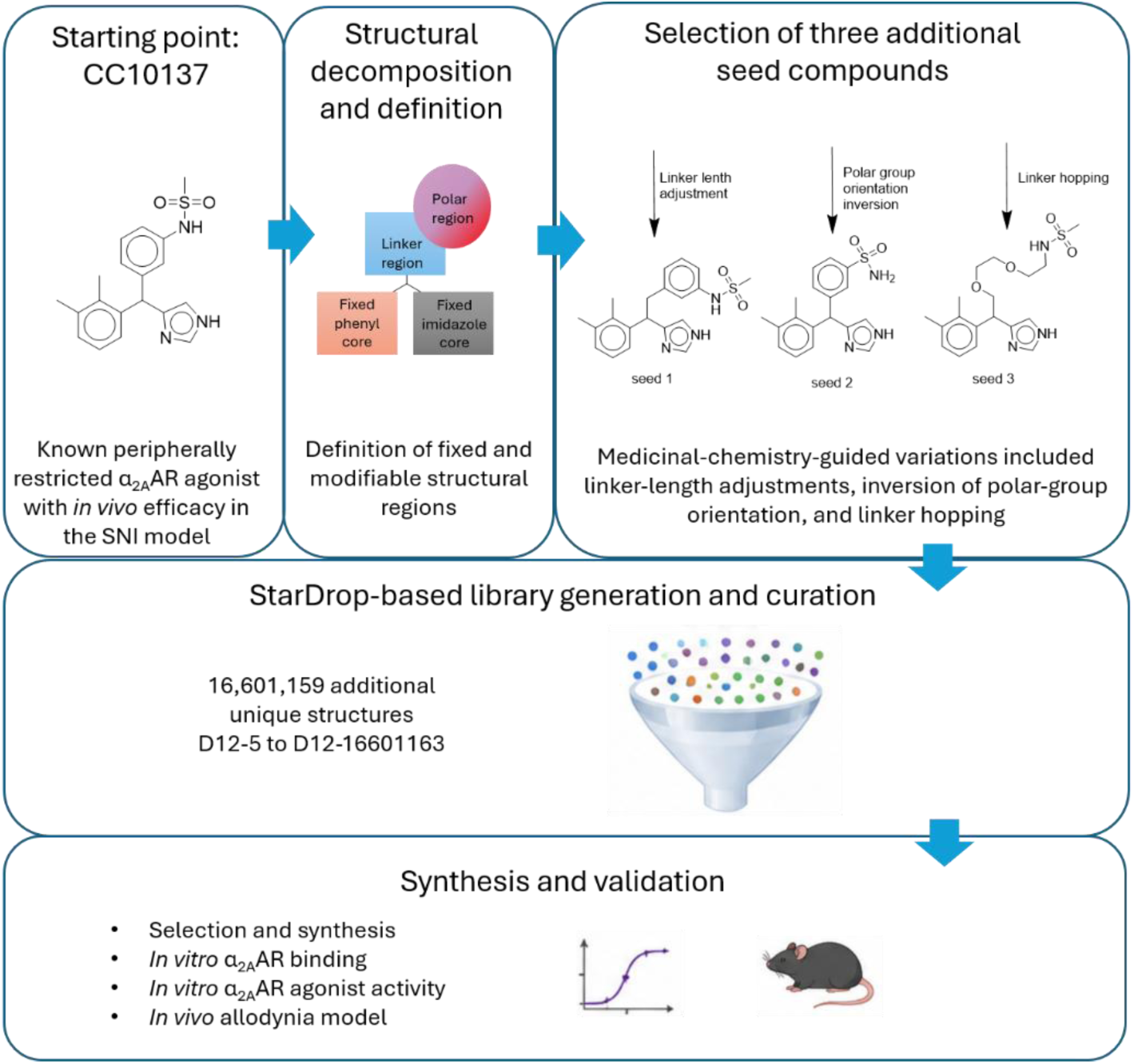
Workflow for CC10137-based chemical-space expansion and experimental evaluation. CC10137 was used as the starting point for structural decomposition into a fixed core and modifiable regions, including linker, polar, and other variable regions. Three additional closely related seed compounds were selected and defined as SAR-informed seed structures based on the earlier medicinal chemistry program, complementing CC10137 to form a set of four seed compounds. These additional seeds incorporated structural variations such as variations in linker length and polar-group orientation, as well as linker hopping. These seed compounds were subjected to StarDrop-based chemical transformation, followed by removal of duplicate structures to generate the nonredundant D12 derivative library. Representative structures from the generated chemical space were subsequently selected for pharmacological characterization, including α_2A_AR binding and agonist activity, and for *in vivo* evaluation in the spared nerve injury (SNI) model.

StarDrop was then used to systematically apply predefined chemical transformation rules to the modifiable regions of the four seed compounds while preserving the fixed receptor-interacting core. After combining the generated structures and removing duplicates, 16,601,159 additional unique structures were obtained and sequentially designated D12-5 through D12-16,601,163. Together with CC10137 (D12-1) and 3 additional seed compounds (D12-2 to D12-4), these structures constituted the D12 derivative library, which was subsequently used for structural comparison and pharmacological evaluation.

The complete α_2A_AR agonist activity concentration-response curves for D12-seed-1 through D12-seed-3 are shown in Figure 2. Together, these results confirmed that the seed set used for chemical-space expansion was anchored by previously reported, experimentally validated α_2A_AR agonists representing distinct SAR-supported structural configurations.

**Figure 2.**
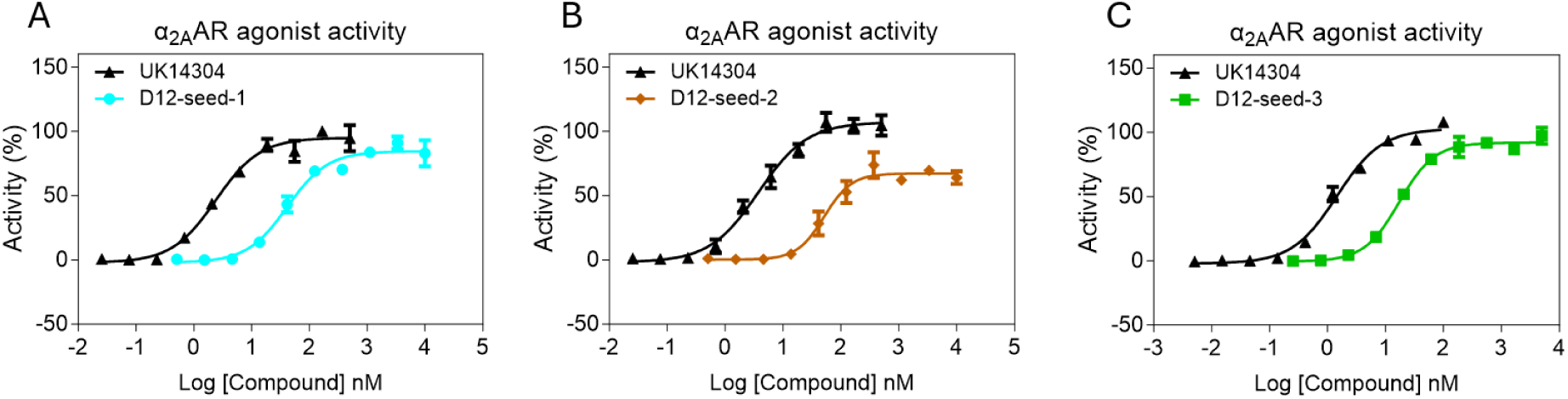
α_2A_AR agonist activity EC_50_ of 3 seed compounds. Concentration-response curves for α_2A_AR activation by (A) D12-seed-1 (EC_50_ = 51.51 nM), (B) D12-seed-2 (EC_50_ = 41.23 nM), (C) D12-seed-3 (EC_50_ = 17.24 nM). Agonist activity was assessed using a calcium flux assay in HEK293 cells stably expressing human α_2A_AR and Gα15. UK14304 was included as the positive reference agonist. Data are presented as mean ± SD, and EC_50_ values were determined by nonlinear regression using the “log(agonist) vs. response – variable slope” model.

### Comparison with previously disclosed compounds

The expanded D12 library, comprising 16,601,163 unique structures, was compared with a defined subset of compounds from our previous medicinal chemistry program focused on modification of the dexmedetomidine-related chiral carbon region. (*8*) This comparison set included the complete Series B (11 compounds), the complete Series D (24 compounds), and Series E-d (8 compounds), comprising all dexmedetomidine-related compounds from Series E, for a total of 43 experimentally characterized structures. These 43 compounds correspond to approximately 0.00026% of the 16,601,163-member D12 library, illustrating the very small fraction of the computationally accessible space represented by the earlier experimentally explored set. Notably, the D12 library generated from only four seed compounds was able to recover all 43 compounds as exact structural matches. This complete recovery reflects both the medicinal chemistry principles encoded in the seed structures and the breadth of chemical-space coverage achieved through StarDrop-based transformation. Thus, a limited set of four carefully selected seed compounds was sufficient to recapitulate the entire set of previously designed analogues within these structural series, while expanding the accessible space to approximately 16.6 million unique structures. These results demonstrate that the combination of SAR-informed seed selection and systematic computational transformation can efficiently reproduce known medicinal chemistry solutions while extending exploration across a substantially broader region of chemical space. To further examine the contribution of individual seed structures, the generated chemical spaces were analyzed according to their seed of origin. The four seed-derived spaces collectively achieved complete recovery of the predefined subset of compounds from the previous medicinal chemistry program. Importantly, the four seeds also generated largely complementary chemical spaces. Of the 16,601,163 unique structures in the final D12 library, the majority were uniquely accessible from a single seed, whereas only a relatively small fraction was shared across multiple seed-derived spaces. Seed-1 and Seed-3 incorporated alternative linker configurations connecting the dexmedetomidine-related region to the polar moiety, whereas Seed-2 represented an inversion of polar-group orientation. These seed configurations were selected based on medicinal chemistry principles and SAR patterns established during the earlier program, and this prior knowledge was important for guiding recovery of previously explored structural space. These results indicate that the use of multiple SAR-informed seeds substantially broadened chemical-space coverage while introducing limited redundancy. A detailed analysis of structural recovery and the contribution of individual seed-derived chemical spaces is shown in Figure 3.

**Figure 3.**
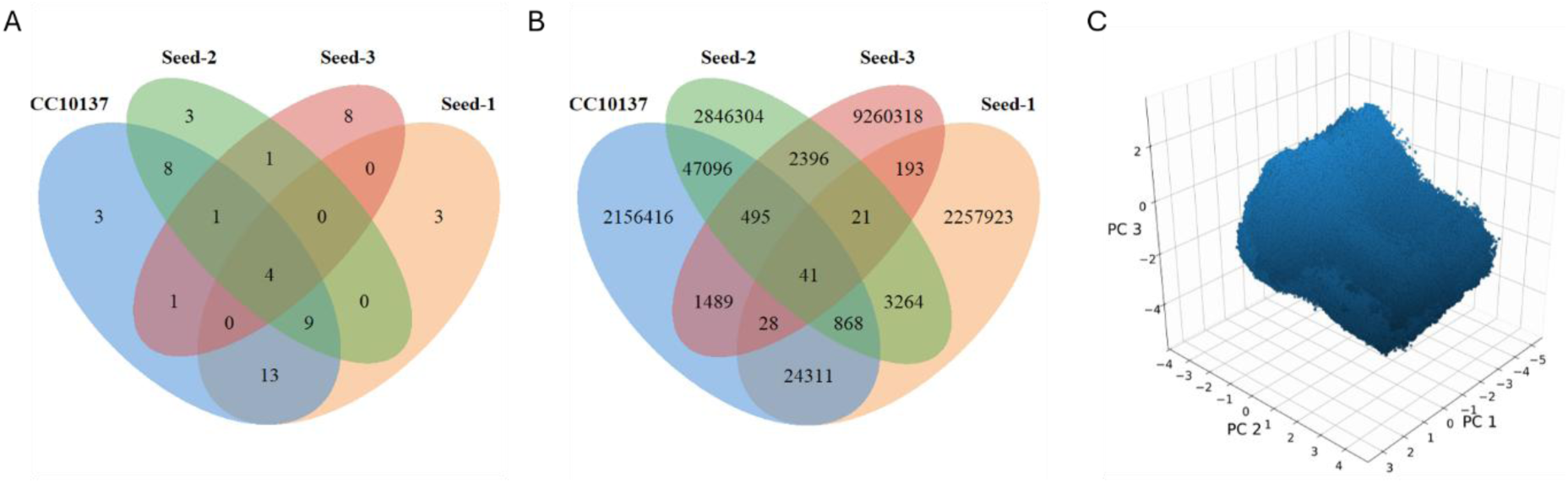
Structural recovery and chemical-space characterization of the four-seed enumeration strategy. (A) Venn analysis showing recovery of the predefined subset of previously characterized compounds by chemical spaces generated from CC10137, D12-seed-1, D12-seed-2, and D12-seed-3. The four seed-derived spaces collectively achieved complete recovery of the comparison set. Numbers indicate compounds uniquely or jointly represented within each seed-derived space. (B) Venn analysis of the complete D12 derivative library showing the distribution of 16,601,163 unique structures generated from the four seed compounds. Numbers indicate structures uniquely generated from each seed or shared among two or more seed-derived spaces. The limited overlap demonstrates substantial complementarity among the four starting structures. (C) Three-dimensional PCA representation of the 16.6 M compound library based on 2048-bit Morgan fingerprints (radius = 2). PC1, PC2, and PC3 accounted for 8.38%, 3.69%, and 2.73% of the total fingerprint variance, respectively, corresponding to a cumulative explained variance of 14.80%. The low cumulative variance reflects the highly multidimensional nature of the generated chemical space.

The complete recovery of these previously reported compounds indicates that the generation rules, together with the SAR information encoded in the four seed structures, were able to systematically sample regions of chemical space that had been independently identified through medicinal chemistry as synthetically accessible and pharmacologically relevant. This finding supports the chemical relevance and coverage of the selected expansion strategy and provides a foundation for applying the workflow prospectively to the identification and prioritization of new analogues.

### PCA characterization of the expanded chemical space

Principal component analysis (PCA) was carried out to give a visual and chemical representation of the chemical space covered by the 16.6 M compound library. The analysis was carried out using 2048-bit Morgan fingerprints (radius=2) with principal component 1 (PC1), PC2, and PC3 accounting for 8.38%, 3.69%, and 2.73% of the total fingerprint variance. The cumulative variance of 14.80% of the first three PCs indicates that the 16.6M compound library is highly multidimensional and cannot be fully represented by the three-dimensional PCA projection. An analysis of the Morgan fingerprint drivers was conducted to identify the chemical and structural features contributing to each PC. PC1 primarily captures variation between the substituted aromatic/heteroaromatic scaffolds and increasingly oxygenated, heteroatom-rich, and side-chain functionalized derivatives. The variation in PC2 distinguishes between the composition of the peripheral functional groups with the positive direction enriched with sulfonyl and nitrogen containing polar groups and the negative direction enriched in halogenated and oxygenated peripheral groups. PC3 captures variation between more functionally substituted analogs and more compact aromatic scaffolds.

### Synthesis and quality control of selected compounds

For further experimental evaluation, five compounds were selected. Two compounds, D12-6223 (previously designated D14(*8*)) from Series D and D12-884990 (previously designated E8(*8*)) from Series E, were selected from the previously reported series as representative structures recovered by the computational workflow. In addition, three compounds, D12-992047, D12-1129389, and D12-2386724, were selected directly from the D12 derivative library to represent distinct structural modification strategies within the generated chemical space. D12-992047 represented a design incorporating an amide-linked oxygen-containing heterocyclic polar group on the aromatic ring; D12-1129389 represented a more flexible polar side chain introduced through an ether/alkyl linker; and D12-2386724 represented a more pronounced scaffold/linker diversification through incorporation of a fused lactam-containing ring system. The structures are shown in Figure 4. D12-992047, D12-1129389, and D12-2386724 were synthesized or resynthesized, and D12-1129389 was further resolved to afford the D12-1129389-I and D12-1129389-II enantiomers. Compound identity was confirmed by ^1^H NMR and LC–MS. Detailed synthetic procedures, analytical results, and quality-control data are provided in the Supporting Information.

**Figure 4.**
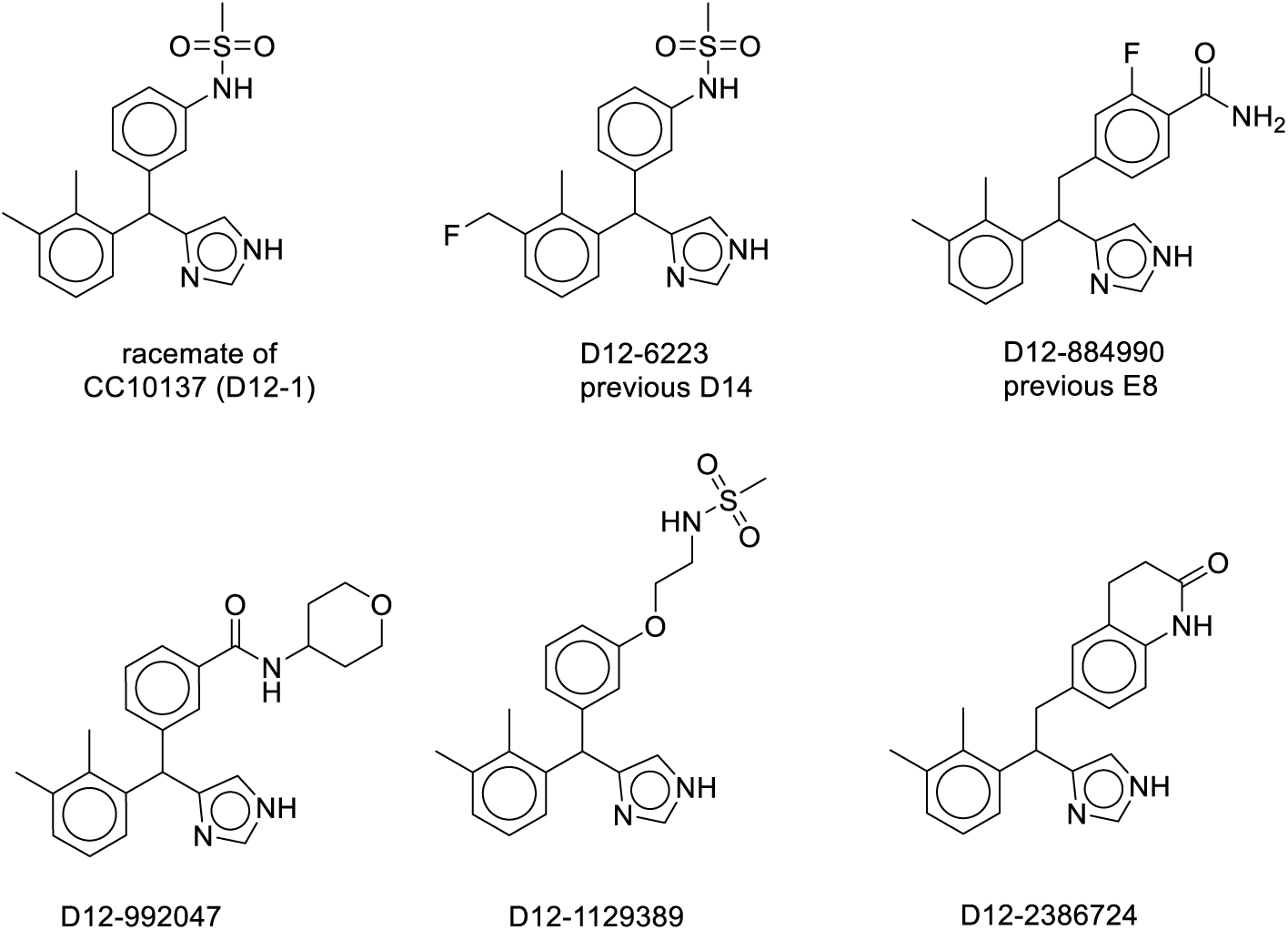
Representative structures from the CC10137-derived chemical-space expansion (D12 derivative library). Representative structures from the D12 derivative library are shown, including the parent compound CC10137 (D12-1), two previously reported compounds recovered by the workflow, D12-6223 (previously D14) and D12-884990 (previously E8), and three additional compounds selected from the generated library, D12-992047, D12-1129389, and D12-2386724. These examples illustrate structural diversity introduced through systematic modification of the phenyl-imidazole linker, peripheral distribution-related substituents, and other predefined variable regions.

### *In vitro* pharmacological evaluation of selected compounds

D12-992047, D12-1129389, and D12-2386724 were first evaluated for α_2A_AR binding affinity, with measured Ki values of 10.15, 23.73, and 2.09 nM, respectively. Chiral resolution of D12-1129389 afforded D12-1129389-I and D12-1129389-II, which showed markedly different Ki of 1036.92 and 2.85 nM, respectively, indicating pronounced stereoselectivity. Binding curves for the α_2A_AR antagonist yohimbine, used as the positive control, together with those for the previously disclosed compounds D12-6223 (23.90 nM) and D12-884990 (5.06 nM), are shown in Figure 5.

**Figure 5.**
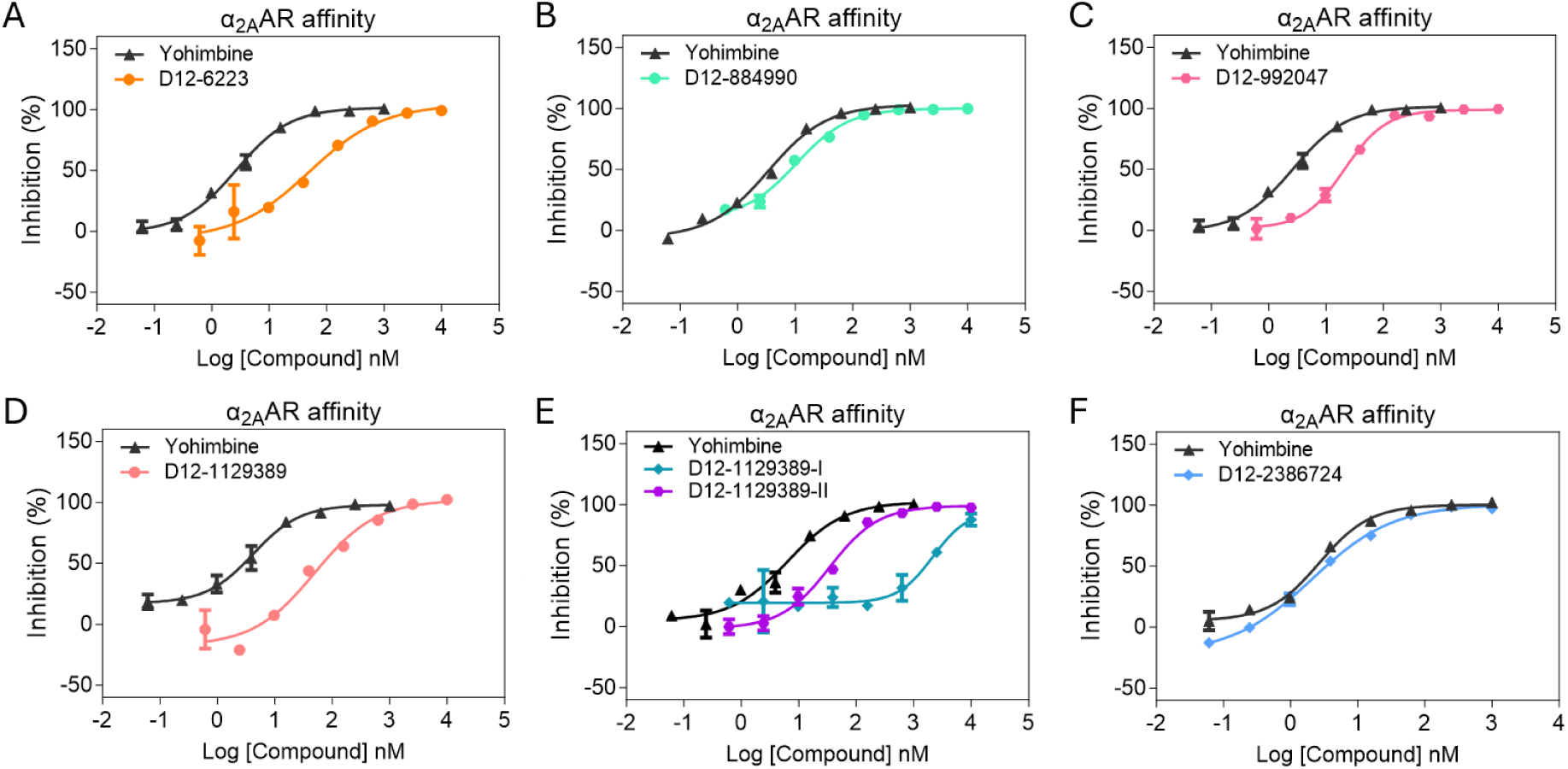
α_2A_AR binding affinities of representative compounds. Concentration-response curves for inhibition of [³H]RX821002 binding to human α_2A_AR by (A) D12-6223 (Ki = 23.90 nM), (B) D12-884990 (Ki = 5.06 nM), (C) D12-992047 (Ki = 10.15 nM), (D) D12-1129389 (Ki = 23.73 nM), (E) D12-1129389-I (Ki = 1036.92 nM) and D12-1129389-II (Ki = 2.85 nM), and (F) D12-2386724 (Ki = 2.09 nM). Yohimbine was included as the positive reference compound in each assay. Data are presented as mean ± SD, and Ki values were determined by nonlinear regression using the “log(inhibitor) vs. response – variable slope” model.

Then, D12-992047, D12-1129389, D12-1129389-II, and D12-2386724, were evaluated for α_2A_AR agonist activity. The corresponding EC_50_ values were 6.432, 15.84, 8.096, and 14.33 nM, respectively. Concentration-response curves for these compounds, together with those for the positive reference agonist UK14304 and the previously disclosed compounds D12-6223 (EC_50_ = 27.95 nM) and D12-884990 (EC_50_ = 6.342 nM), are shown in Figure 6. Overall, the newly evaluated compounds retained sub-25 nM α_2A_AR binding affinity and agonist potency. These findings provide experimental support that the computational workflow sampled chemically relevant structures compatible with α_2A_AR binding and activation.

**Figure 6.**
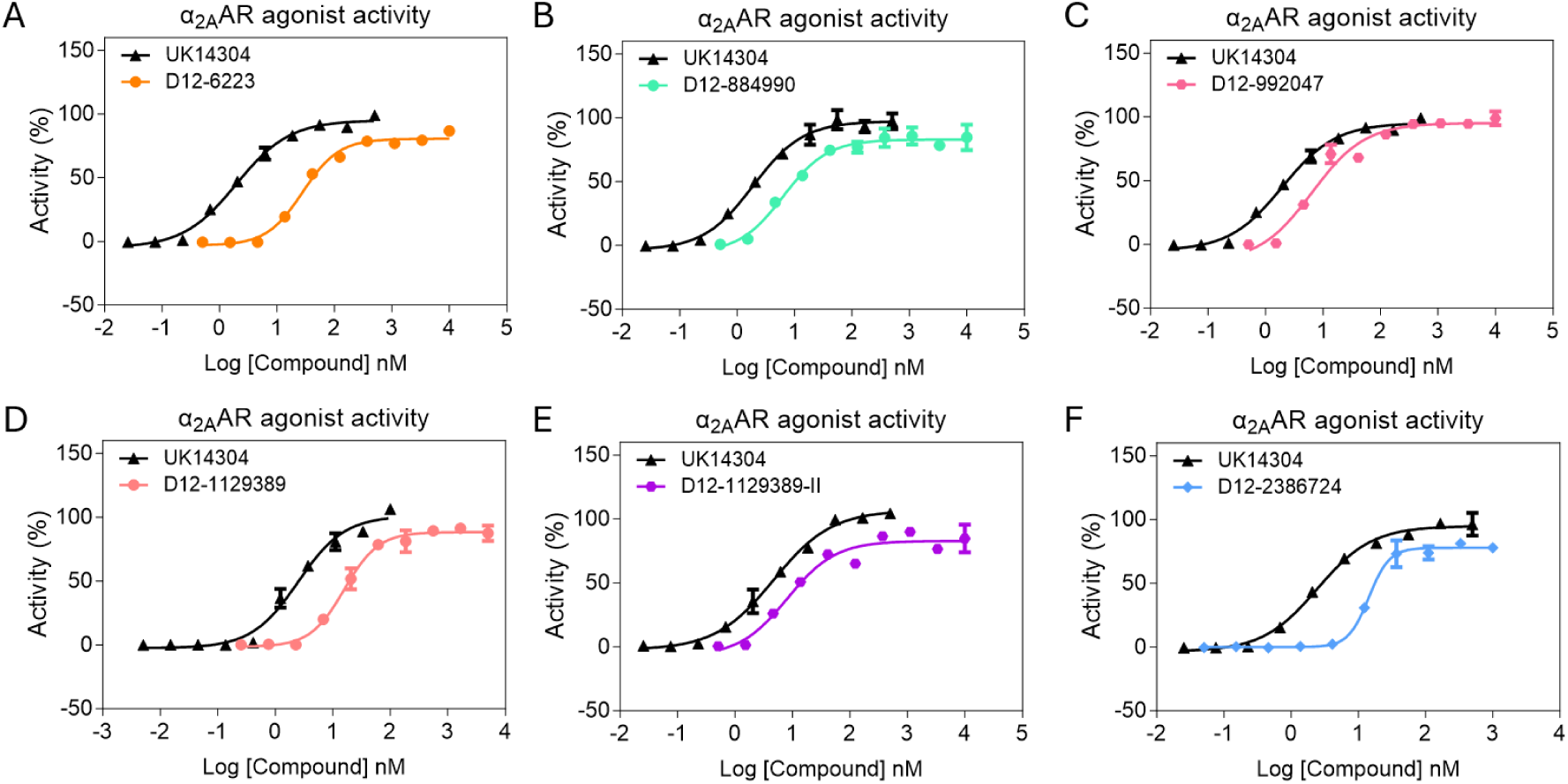
α_2A_AR agonist activities of representative compounds. Concentration-response curves for α_2A_AR activation by (A) D12-6223 (EC_50_ = 27.95 nM), (B) D12-884990 (EC_50_ = 6.342 nM), (C) D12-992047 (EC_50_ = 6.432 nM), (D) D12-1129389 (EC_50_ = 15.84 nM), (E) D12-1129389-II (EC_50_ = 8.096 nM), and (F) D12-2386724 (EC_50_ = 14.33 nM). Agonist activity was assessed using a calcium flux assay in HEK293 cells stably expressing human α_2A_AR and Gα15. UK14304 was included as the positive reference agonist. Data are presented as mean ± SD, and EC_50_ values were determined by nonlinear regression using the “log(agonist) vs. response – variable slope” model.

### *In vivo* evaluation in the SNI model

D12-6223 (2 mg/kg), D12-884990 (2 mg/kg), D12-992047 (2 mg/kg), D12-1129389 -II (1 mg/kg), and D12-2386724 (2 mg/kg), were evaluated in the spared nerve injury (SNI) model shown in Figure 7. All five compounds produced significant anti-allodynic effects, with inhibition rates of 39.3%, 55.7%, 42.0%, 46.1%, and 41.0%, respectively. The consistent retention of *in vivo* activity across structurally distinct compounds indicates that meaningful structural variation within the generated chemical space can be accommodated without loss of pharmacologically relevant efficacy. Importantly, these results extend the validation of the computational workflow beyond structural similarity and *in vitro* receptor activity, demonstrating that compounds sampled from the expanded chemical space can also retain functional efficacy in an established neuropathic pain model.

**Figure 7.**
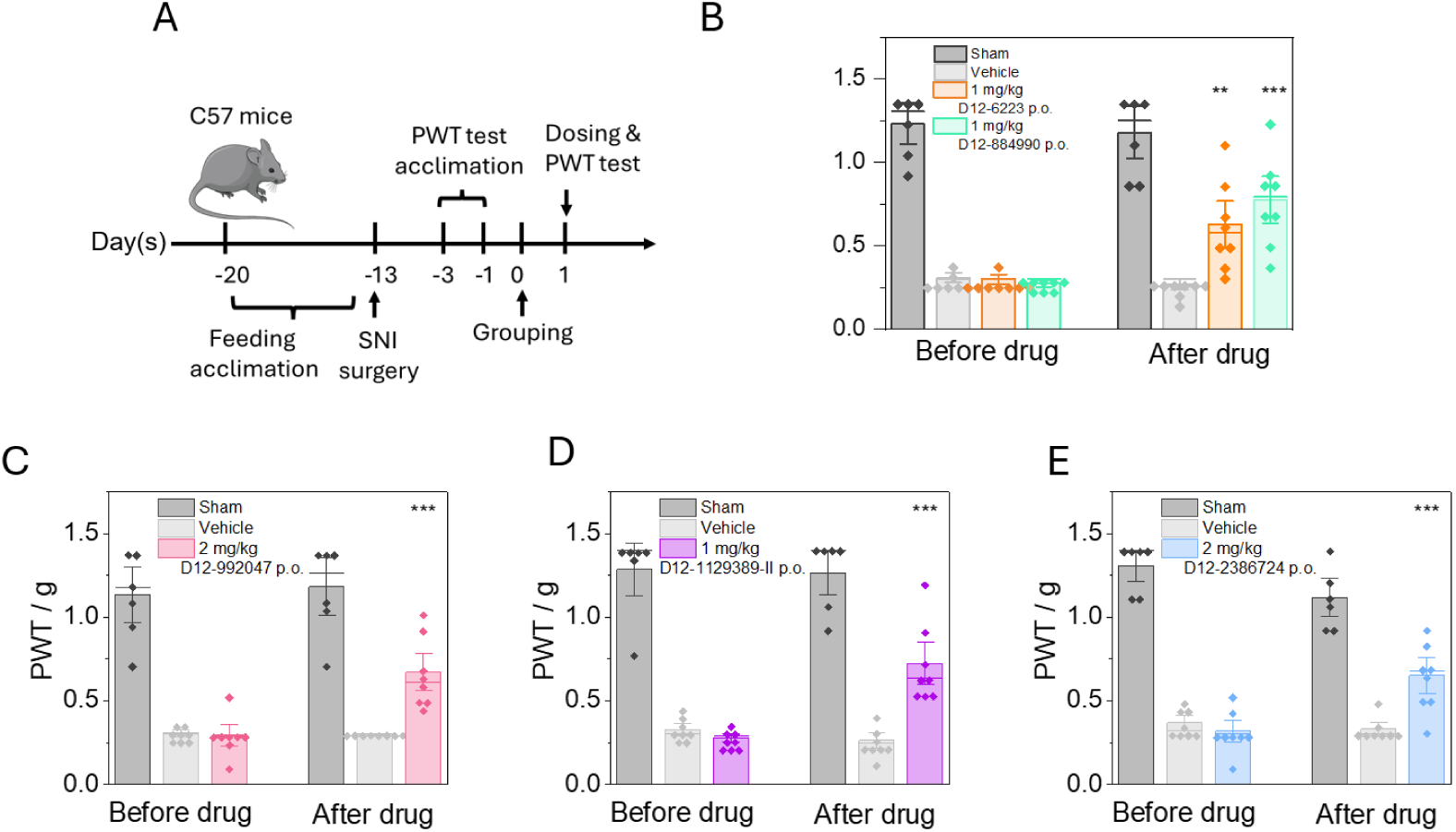
Anti-allodynic effects of representative compounds in the spared nerve injury model. (A) Schematic illustration of the spared nerve injury (SNI) model and experimental design. On day 13 after surgery, eligible mice were randomized into treatment groups and orally administered the indicated test compounds. (B-E) Paw withdrawal threshold (PWT) measured at baseline and 1 h after oral administration of D12-6223 (2 mg/kg), D12-884990 (2 mg/kg), D12-992047 (2 mg/kg), D12-1129389-II (1 mg/kg), and D12-2386724 (2 mg/kg), respectively. Sham-operated and vehicle-treated mice were included as controls. The corresponding inhibition rates at 1 h were 39.3%, 55.7%, 42.0%, 46.1%, and 41.0%, respectively. Data are presented as mean ± SEM. Statistical significance between pre-dose and post-dose measurements was determined using a paired t-test. The p values for the five compounds were 0.00548 (**), 0.000543 (***), 0.000132 (***), 0.000370 (***), and 0.000525 (***), respectively. **p < 0.01; ***p < 0.001.

Together, these data show that selected structures sampled from the generated chemical space can retain both α_2A_AR pharmacological activity and *in vivo* anti-allodynic efficacy. The ability to recover both previously validated compounds and additional structures selected from the generated library with retained *in vitro* and *in vivo* activity supports the use of this workflow not only as a retrospective tool for reproducing medicinal chemistry outcomes, but also as a prospective strategy for prioritizing new analogues for further optimization and experimental evaluation.

## Discussion

The present study shows that a small number of SAR-informed seed structures can support large-scale and structurally broad exploration of the chemical space surrounding a first-in-class lead. Starting from CC10137 and three related seed compounds, systematic rule-based transformation generated approximately 16.6 million unique structures. The resulting library covered extensive variation across the predefined modifiable regions of the scaffold and substantially expanded the structural landscape accessible from the original series.

We next examined how this expanded space related to the chemistry previously explored experimentally. The 43 compounds selected for retrospective comparison represented only approximately 0.00026% of the 16.6-million-member library, yet all 43 were recovered as exact structural matches. These compounds had been independently designed, synthesized, and experimentally characterized in the earlier CC10137 program, providing a direct benchmark for the relevance of the enumerated space. Their complete recovery indicates that the computational library encompassed the experimentally established regions of the series while extending into a much larger surrounding chemical space.

The contribution of the individual seed structures was evident from the Venn analysis. The four seed-derived spaces were highly complementary and showed relatively limited overlap. Each seed therefore contributed access to distinct structural regions, consistent with the differences in linker arrangement, polar-group orientation, and related SAR-supported features encoded in the starting structures. These results emphasize the importance of seed selection in scaffold-based enumeration and show how multiple pharmacologically relevant starting points can broaden the accessible chemical space.

PCA provided an independent view of the structural diversity within the full library. The first three principal components accounted for only 14.80% of the total Morgan fingerprint variance, consistent with a highly multidimensional chemical space. Examination of the fingerprint features contributing to these components revealed variation in aromatic and heteroaromatic substitution, heteroatom and oxygen content, peripheral polar functionality, and the extent of side-chain modification. Together, the Venn and PCA analyses show that the expanded library spans multiple structural dimensions and captures diversity that cannot be represented by a small number of closely related analogue series.

The computationally generated space was then evaluated experimentally. D12-992047, D12-1129389, and D12-2386724, selected directly from the expanded library, retained α_2A_AR binding affinity and agonist potency below 25 nM. Chiral resolution of D12-1129389 further revealed pronounced stereoselectivity, with D12-1129389-II showing substantially greater binding affinity than D12-1129389-I, consistent with a strong dependence of α_2A_AR recognition on stereochemical configuration. *In vivo* evaluation provided an additional level of validation. Five representative compounds, including previously characterized structures recovered by the workflow and compounds selected directly from the expanded library, were tested in the SNI model, and all produced significant anti-allodynic effects, with inhibition rates ranging from 39.3% to 55.7%. These findings establish that the expanded chemical space contains compounds that retain both receptor-level pharmacological activity and *in vivo* efficacy.

This type of systematic exploration is particularly relevant for first-in-class series, where experimentally defined SAR typically covers only a small fraction of the accessible structural landscape. Scaffold-seeded enumeration provides a way to extend experimentally established medicinal chemistry knowledge into a much larger surrounding space while preserving structural features important for activity. In this setting, SAR-informed seed selection connects experimentally derived knowledge with large-scale virtual exploration and provides a framework for identifying additional regions of chemical space for subsequent optimization.

The framework described here is readily extendable beyond the CC10137-derived D12 library. Following completion of the present study, we began applying the same strategy to additional α_2A_AR agonist scaffolds, including other clinically approved agonists, compounds in clinical development, and newly disclosed chemical series. These ongoing efforts are being used to extend the accessible virtual space from the millions of structures examined here toward libraries containing billions of structures. These expanded datasets are not included in the analyses presented in this study. Future extensions could also incorporate isotopic substitution, such as deuteration, to further broaden the scope of molecular design.

The same design logic can also be implemented in other computational environments supporting rule- or reaction-based enumeration, including both commercial platforms and open-source implementations. The core elements remain unchanged: SAR-informed seed selection, definition of conserved and modifiable regions, systematic application of chemically meaningful transformations, and integration of the resulting seed-derived spaces. This platform-independent framework provides a general strategy for mapping the chemical space around α_2A_AR agonist scaffolds.

Overall, this workflow provides a practical approach for translating medicinal chemistry knowledge into systematically generated chemical spaces that can be analyzed, prioritized, synthesized, and experimentally tested. For newly established series with limited experimental coverage, scaffold-seeded enumeration can reveal a much broader structural landscape than is accessible through analogue-by-analogue design and synthesis alone. In the present study, the combination of large-scale enumeration, complete recovery of the predefined experimental comparison set, multidimensional structural characterization, and the *in vitro* and *in vivo* validation supports the use of this strategy for continued exploration of the CC10137 series and broader chemical-space mapping of α_2A_AR agonists.

## Supplemental File

The full dataset comprising 16,601,163 SMILES generated from the transformation of the 4 seed compounds can be found on figshare (10.6084/m9.figshare.33868108).

## Data Resource

Following completion of the present study, we have continued to expand the chemical space using the same computational framework, together with additional structural templates and transformation rules. This ongoing expansion extends beyond the scope of the present study and was not included in the computational or experimental analyses reported here. The resulting expanded molecular datasets are being made publicly available at figshare.com and maker-ip.com.

## Methods

### Compound generation

Compounds were generated using StarDrop (Optibrium Ltd.) and its Chemical Transformation method.(*27*) CC10137 was first structurally decomposed into a fixed receptor-interacting core and several predefined modifiable regions, including the linker, polar, and other variable regions. Based on this framework and analysis of the structure-activity relationships established during the earlier CC10137 program, three closely related structural variants, D12-seed-1 to D12-seed-3, were selected from the earlier CC10137 medicinal chemistry program. Together with CC10137, these compounds formed a set of four SAR-informed seed structures for computational expansion. The three additional seeds incorporated modest but deliberate structural variations, including positional changes, small adjustments in linker length, reversal of the orientation of selected polar functional groups, and linker hopping, with the aim of encoding selected medicinal chemistry and SAR principles into the starting structures before computational expansion.

For each seed compound, the known receptor-interacting core was kept fixed to minimize disruption of α_2A_AR activity, while chemical transformations were directed toward the predefined modifiable regions. Approximately 30,000 precedented chemical transformation rules available in StarDrop were enabled to provide broad coverage of medicinal-chemistry-relevant structural modifications and maximize the diversity of structures generated from the predefined modifiable regions. Structures generated from the four seed compounds were combined, and duplicate structures were removed to generate the final nonredundant D12 derivative library.

For selection of representative compounds for subsequent experimental evaluation, preference was given to structures containing polar functional groups, including amines, amides, and sulfonamides, as well as structural features associated with P-gp-mediated efflux. StarDrop version 8.0.1.9523 was used for these operations.

### PCA analysis

Principal component analysis (PCA) was carried out to give a visual and chemical representation of the chemical space covered by the 16.6 M compound library. To generate the three-dimensional PCA plot every valid molecule was converted into a 2048-bit Morgan fingerprint (radius = 2) using RDKit(*28*). The covariance matrix of the resulting 2048-bit fingerprint was calculated, and the principal components (PCs) were determined following eigendecomposition of the covariance matrix using NumPy(*29*). Each compound was then assigned a coordinate on the 3D plot by projecting its Morgan fingerprint onto PC1, PC2, and PC3 and then rendered with Matplotlib to give a representation of the molecular fingerprint variation across the chemical library(*30, 31*). Finally, the Morgan fingerprint bits with the largest positive and negative loadings for PC1, PC2, and PC3 were identified, and representative molecular substructures associated with these bits were examined to facilitate chemical interpretation of each principal component. The following software was used for the analysis: Python 3.14.3, RDKit 2026.03.1, Numpy 2.4.3, SciPy1.18.1, and Matplotlib 3.11.1.

### Chemical synthesis and quality control

#### D12-992047 (3-((2,3-dimethylphenyl)(1*H*-imidazol-4-yl)methyl)-*N*-(tetrahydro-2*H*-pyran-4-yl)benzamide)

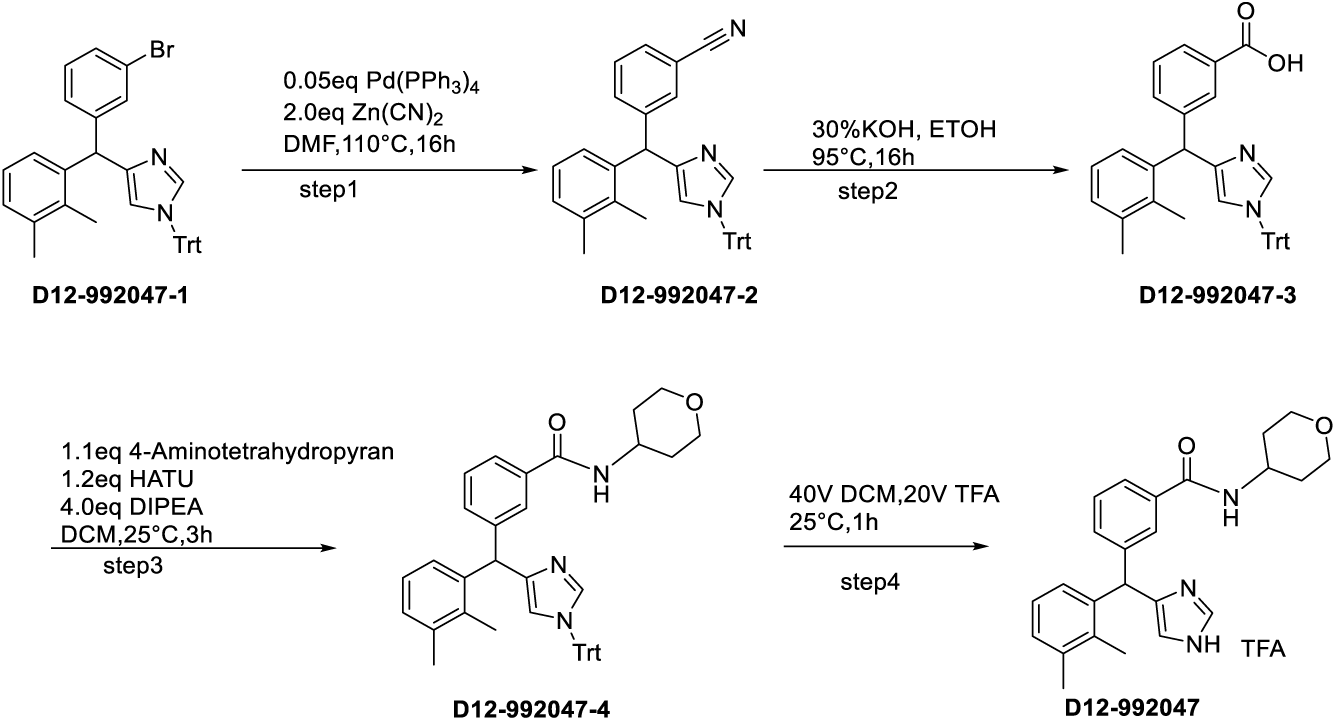

Step 1: A mixture of D12-992047-1 (4-((3-bromophenyl)(2,3-dimethylphenyl)methyl)-1-trityl-1*H*-imidazole, 250 mg, 0.43 mmol, 1.0 eq), Zn(CN)_2_ (100 mg, 0.86 mmol, 2.0 eq), and Pd(PPh_3_)_4_ (25 mg, 0.02 mmol, 0.05 eq) in DMF (2.55 mL) was stirred at 110 °C for 16 h. Upon completion, as confirmed by LC–MS, the reaction mixture was concentrated under reduced pressure. The residue was purified by column chromatography to afford D12-992047-2 (180 mg). Yield: 79.1%.

Step 2: A mixture of D12-992047-2 (180 mg, 0.34 mmol, 1.0 eq) in EtOH (6 mL) and 30% aqueous KOH (2 mL) was stirred at 105 °C for 16 h. Upon completion, as confirmed by LC–MS, the reaction mixture was concentrated under reduced pressure. Water (2 mL) was added, and the mixture was adjusted to pH 3–4 with 4 N HCl. The resulting solid was collected by filtration to afford D12-992047-3 (150 mg). Yield: 79.4%.

Step 3: A mixture of D12-992047-3 (150 mg, 0.27 mmol, 1.0 eq), 4-aminotetrahydropyran (30 mg, 0.30 mmol, 1.1 eq), and DIPEA (142 mg, 1.10 mmol, 4.0 eq) in DCM (3 mL) was cooled to 0 °C. HATU (125 mg, 0.33 mmol, 1.2 eq) was added, and the reaction mixture was stirred at 25 °C for 2 h. Upon completion, as confirmed by LC–MS, the reaction mixture was concentrated under reduced pressure. The residue was purified by column chromatography to afford D12-992047-4 (100 mg). Yield: 59.3%.

Step 4: A solution of D12-992047-4 (100 mg, 0.16 mmol, 1.0 eq) in DCM (4 mL) was cooled to 0 °C, and TFA (2 mL) was added. The reaction mixture was stirred at 25 °C for 1 h. Upon completion, as confirmed by LC–MS, the reaction mixture was concentrated under reduced pressure. The residue was purified by preparative HPLC to afford D12-992047 (30 mg, purity: 99%) as a white powder. Yield: 37.2%.

LC–MS: [M+H]^+^ = 390.2.

^1^H NMR (400 MHz, DMSO+D2O) δ 9.02 (d, J = 1.1 Hz, 1H), 7.78 (d, J = 7.8 Hz, 1H), 7.67 (s, 1H), 7.45 (t, J = 7.7 Hz, 1H), 7.28 (d, J = 7.7 Hz, 1H), 7.16 – 7.06 (m, 2H), 6.93 (s, 1H), 6.67 (d, J = 7.4 Hz, 1H), 5.89 (s, 1H), 3.92 (ddd, J = 29.2, 18.1, 8.0 Hz, 3H), 3.36 (t, J = 11.1 Hz, 2H), 2.24 (s, 3H), 2.11 (s, 3H), 1.75 – 1.68 (m, 2H), 1.56 (qd, J = 11.9, 4.2 Hz, 2H).

#### D12-1129389: *N*-(2-(3-((2,3-dimethylphenyl)(1*H*-imidazol-4-yl)methyl)phenoxy)ethyl)methanesulfonamide

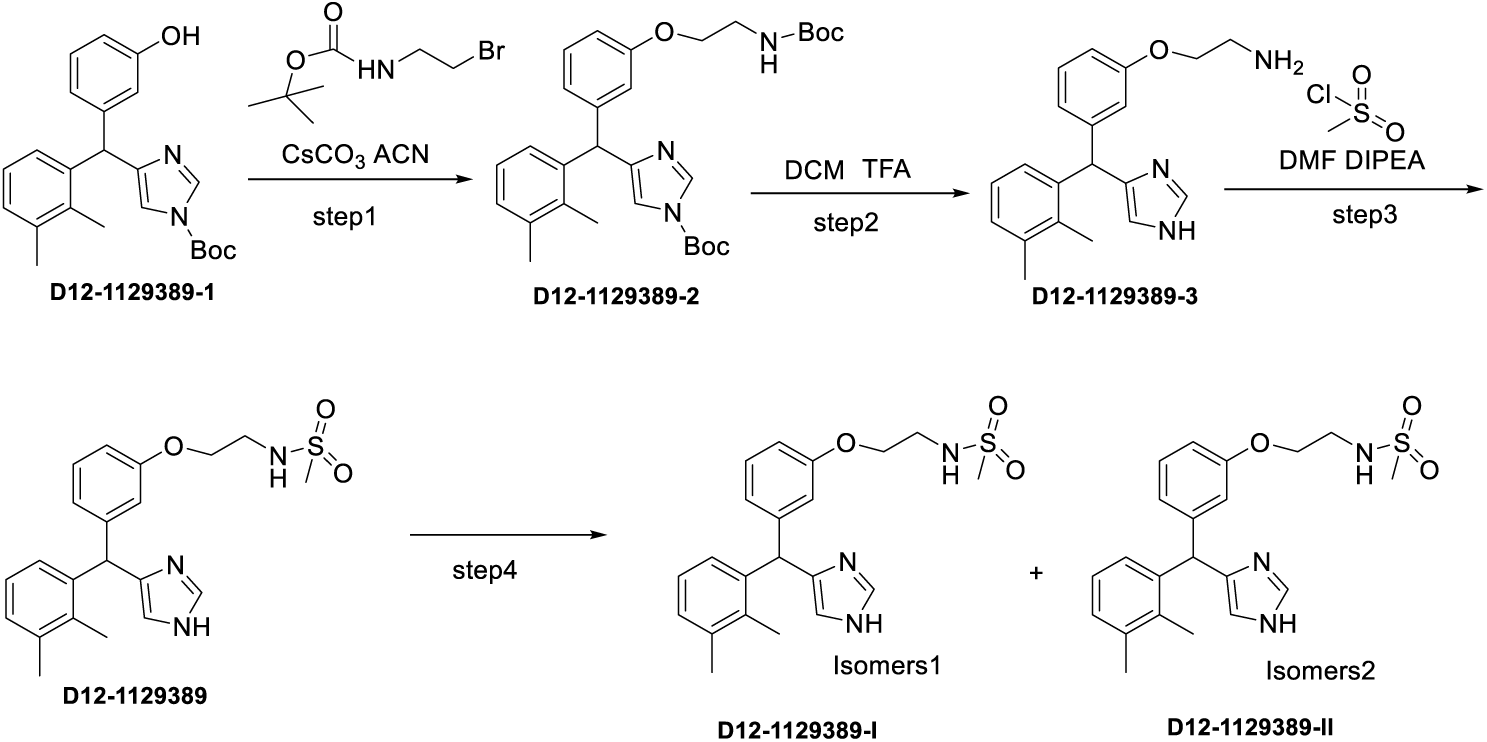

Step 1: To a 100 mL three-necked flask were added D12-1129389-1 (*tert*-butyl 4-((2,3-dimethylphenyl)(3-hydroxyphenyl)methyl)-1*H*-imidazole-1-carboxylate, 3.5 g, 9.25 mmol, 1 eq), 39684-80-5 (4.2 g, 18.49 mmol, 2 eq), Cs_2_CO_3_ (4.5 g, 13.87 mmol, 1.5 eq), and ACN (40 mL). The reaction mixture was stirred at 60 °C for 12 h. Upon completion, as confirmed by LCMS, the reaction mixture was poured into water and extracted with EtOAc. The organic phase was dried over Na_2_SO_4_ and concentrated under reduced pressure to afford D12-1129389-2 (2.3 g) as a white solid, which was used directly in the next step without further purification. Yield: 47.7%.

Step 2: To a 100 mL three-necked flask were added D12-1129389-2 (2.3 g), DCM (30 mL), and TFA (15 mL). The reaction mixture was stirred at 25 °C for 12 h. Upon completion, as confirmed by LCMS, the reaction mixture was poured into water and adjusted to pH 10, followed by extraction with DCM. The organic phase was dried over Na₂SO₄ and concentrated under reduced pressure. The residue was purified by silica gel column chromatography to afford D12-1129389-3 (1.0 g) as a yellow solid. Yield: 70.5%.

Step 3: To a 10 mL three-necked flask were added D12-1129389-3 (900 mg, 2.80 mmol, 1 eq), DIPEA (905 mg, 7.0 mmol, 2.5 eq), methanesulfonyl chloride (353 mg, 3.08 mmol, 1.1 eq), and DMF (5 mL). The reaction mixture was stirred at 25 °C for 2 h. Upon completion, as confirmed by LCMS, the reaction mixture was poured into water and extracted with EtOAc. The organic phase was dried over Na_2_SO_4_ and concentrated under reduced pressure. The residue was purified by silica gel column chromatography to afford D12-1129389 (420 mg) as a white solid. Yield: 37.5%.

Step 4: D12-1129389 (420 mg) was separated by preparative chiral HPLC (column: CHIRALPAK® IG, 10 µm, 30 × 250 mm; mobile phase A: hexane containing 0.2% DEA; mobile phase B: EtOH containing 0.2% DEA; flow rate: 25 mL/min) to afford D12-1129389-I (166 mg, purity: 98.3%, yield: 79%), the first-eluting enantiomer, as a white powder, and D12-1129389-II (170 mg, purity: 99.5%, yield: 81%), the second-eluting enantiomer, as a white powder.

D12-1129389-I: LC–MS: [M+H]^+^=400.2

1H NMR (400 MHz, DMSO+D2O) δ 7.58 (d, J = 0.9 Hz, 1H), 7.20 (t, J = 7.7 Hz, 1H), 7.11 – 6.86 (m, 2H), 6.74 (dd, J = 26.0, 7.5 Hz, 3H), 6.64 (s, 1H), 6.44 (s, 1H), 5.56 (s, 1H), 3.93 (t, J = 5.3 Hz, 2H), 3.27 (t, J = 5.3 Hz, 2H), 2.89 (s, 3H), 2.20 (s, 3H), 2.09 (s, 3H). D12-1129389-II: LC–MS: [M+H]^+^=400.2 1H NMR (400 MHz, DMSO+D2O) δ 7.58 (s, 1H), 7.20 (t, J = 7.7 Hz, 1H), 6.98 (d, J = 8.5 Hz, 2H), 6.73 (dd, J = 26.9, 7.4 Hz, 3H), 6.62 (s, 1H), 6.44 (s, 1H), 5.56 (s, 1H), 3.26 (t, J = 5.2 Hz, 2H), 2.89 (s, 3H), 2.19 (s, 3H), 2.08 (s, 3H).

#### D12-2386724: 6-(2-(2,3-dimethylphenyl)-2-(1*H*-imidazol-4-yl)ethyl)-3,4-dihydroquinolin-2(1*H*)-one

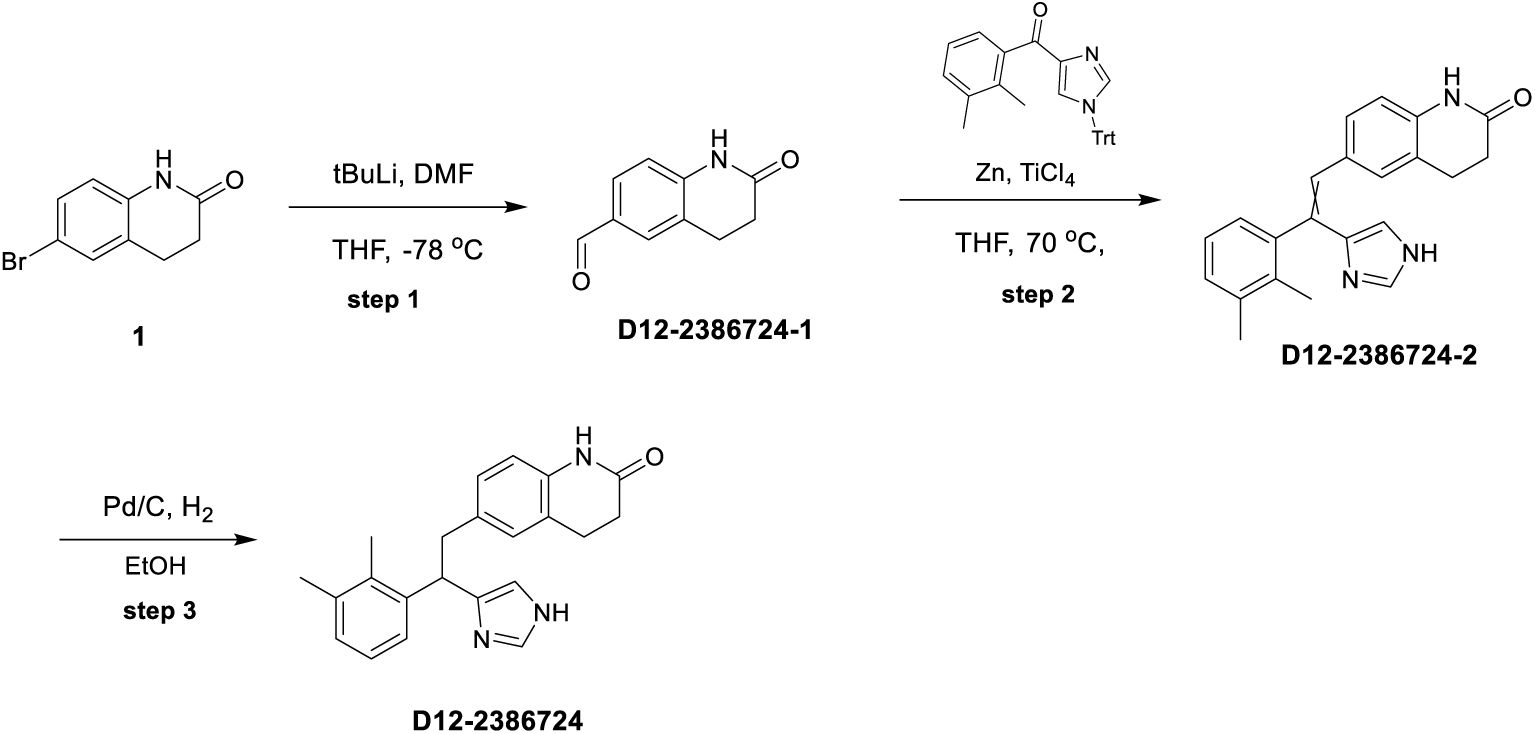

Step 1: A solution of 6-bromo-3,4-dihydro-1*H*-quinolin-2-one (CAS No. 3279-90-1; 3.00 g, 13.3 mmol, 1.0 eq) in THF (30 mL) was cooled to -78 °C under a nitrogen atmosphere. *n*-BuLi (2.5 M in hexane, 21.3 mL, 53.2 mmol, 4.0 eq) was added, and the reaction mixture was stirred at -78 °C for 0.5 h. *N,N*-Dimethylformamide (1.84 g, 15.9 mmol, 1.2 eq) was then added, and the reaction mixture was stirred at −78 °C for an additional 3 h. Upon completion, as confirmed by LC–MS, the reaction mixture was quenched with aqueous NH_4_Cl and diluted with EtOAc (500 mL). The organic phase was washed with water (100 mL × 3), concentrated under reduced pressure, and purified by flash column chromatography (EtOAc/hexanes, 0–50%) to afford D12-2386724-1 (1.20 g) as a white solid. Yield: 41.3%.

LC–MS: [M+H]⁺ = 176.1.

Step 2: A mixture of 4-[(2,3-dimethylphenyl)carbonyl]-1-(triphenylmethyl)imidazole (694.7 mg, 1.57 mmol, 1.1 eq), D12-2386724-1 (250 mg, 1.43 mmol, 1.0 eq), and Zn powder (560.1 mg, 8.56 mmol, 6.0 eq) in THF (15 mL) was cooled to -10 °C under a nitrogen atmosphere. TiCl_4_ (1.62 g, 8.56 mmol, 6.0 eq) was added dropwise, and the reaction mixture was stirred at -10 °C for 0.5 h. The mixture was then heated to 70 °C and stirred for 4 h. After cooling to room temperature, the reaction mixture was poured into water (100 mL) and extracted with EtOAc (200 mL). The organic phase was washed with water, dried over Na_2_SO_4_, and concentrated under reduced pressure. The residue was purified by silica gel column chromatography (MeOH/DCM, 0-5%) to afford D12-2386724-2 (250 mg) as a yellow solid. Yield: 45.7%.

LC–MS: [M+H]⁺ = 344.1.

Step 3: A mixture of D12-2386724-2 (250 mg, 0.757 mmol, 1.0 eq), Pd/C (32.2 mg), and Pd(OH)_2_ (42.5 mg) in EtOH (5 mL) was stirred at 25 °C for 12 h under a hydrogen atmosphere. Upon completion, the reaction mixture was filtered and concentrated under reduced pressure. The crude product was purified by preparative HPLC (instrument: SHIMADZU LC-20AP-4; column: Gemini; mobile phase: ACN/H₂O containing 0.1% TFA; gradient: 0-30% ACN) to afford D12-2386724 (58.7 mg) as a white solid. Yield: 22.4%.

LC–MS: [M+H]⁺ = 346.15.

^1^H NMR (400 MHz, DMSO) δ 14.13 (s, 1H), 9.99 (s, 1H), 8.95 (s, 1H), 7.62 (s, 1H), 7.15–7.05 (m, 3H), 6.96–6.88 (m, 2H), 6.66 (d, *J* = 4.0 Hz, 1H), 4.64 (t, *J* = 6.0 Hz, 1H), 3.26–3.20 (m, 1H), 3.03 (m, 1H), 2.79–2.73 (m, 2H), 2.41–2.36 (m, 2H), 2.22 (s, 3H), 2.14 (s, 3H).

Overall yield: 4.23%.

### In vitro pharmacology: α_2A_AR agonist activity

The study was conducted at WuXi AppTec according to its standardized *in vitro* pharmacology protocol for GPCR calcium flux assays. HEK293 cells stably expressing human α_2A_AR and Gα15 were provided by WuXi AppTec and used to evaluate agonist activity. Cells were maintained in DMEM supplemented with 10% FBS, 1% penicillin-streptomycin, 300 µg/mL G418, and 2 µg/mL BSA. The stable cell line had been generated by transfection followed by selection with geneticin and blasticidin S to obtain a single clone expressing α_2A_AR, which was subsequently used for calcium flux and binding assays.

For the calcium flux assay, cells were washed once with DPBS (21-0310CVR, Corning) and detached with 0.05% trypsin-EDTA (25300-062, Gibco) at 37 °C for 1 min. Trypsinization was terminated by addition of culture medium, and the cell suspension was centrifuged at 1,000 rpm for 5 min. The supernatant was discarded, and the cell pellet was resuspended in fresh culture medium. Cell number and viability were determined using a Vi-CELL™ analyzer (Beckman), and the cell density was adjusted to 1 × 10^6^ cells/mL. A 20 µL aliquot of the cell suspension, corresponding to 20,000 cells per well, was seeded into 384-well plates (Greiner) and incubated overnight at 37 °C in 5% CO₂.

On the following day, a 2× Fluo-4 Direct dye solution (8 µM; Invitrogen) was prepared in 10 mL of FLIPR buffer consisting of 20 mM HEPES (Gibco), 1× HBSS, and 0.5% BSA, supplemented with 0.2 mL of 250 mM probenecid, and mixed for 5 min. The culture medium was aspirated, and 20 µL of the 2× Fluo-4 reagent was added to each well. Plates were incubated at 37 °C for 50 min and then equilibrated at room temperature for 10 min.

Test compounds were serially diluted threefold in DMSO to generate ten concentrations. DMSO served as the low-signal control. A 30 µL aliquot of assay buffer was added to each well of the cell plate, followed by centrifugation at 1,000 rpm, and 10 µL of compound solution was then added to each well. Fluorescence signals were recorded in real time using a FLIPR Tetra+ system (Molecular Devices). Activation (%) was calculated as (RFU_sample − RFU_DMSO)/(RFU_HC − RFU_DMSO) × 100, where RFU represents relative fluorescence units averaged over readings 1-90, and HC represents the highest concentration of the positive reference compound, UK14304. EC_50_ values were determined using the “log(agonist) vs. response − variable slope” model in GraphPad Prism 5.0.

### In vitro pharmacology: α_2A_AR radioligand binding experiment

The assay was conducted at WuXi AppTec according to its standardized *in vitro* pharmacology protocol for α_2A_AR binding assessment. Membranes containing human α_2A_AR (NM_000681) were prepared from a stable α_2A_AR-expressing cell line. Test compounds and the reference compound yohimbine were serially diluted fourfold to generate eight concentrations, with maximum starting concentrations of 10 µM and 1,000 nM, respectively. For each assay, 1 µL of test compound solution was transferred to the assay plate. For the nonspecific binding control (low control, LC) and total binding control (high control, HC), 1 µL of 0.2 mM yohimbine and 1 µL of DMSO were added, respectively.

Each sample was then mixed with 100 µL of membrane suspension containing 5 µg of membrane protein and [³H]RX821002 (0.5 nM). The assay and wash buffer consisted of 50 mM Tris-HCl, pH 7.4. Binding reactions were incubated at room temperature for 1 h. In parallel, Unifilter-96 GF/C filter plates (6005174, PerkinElmer) were pretreated with 50 µL of 0.3% polyethyleneimine (PEI) for at least 0.5 h at room temperature. The binding mixtures were filtered through the GF/C plates using a PerkinElmer UNIFILTER-96 Cell Harvester and washed four times with cold assay buffer. After drying at 50 °C for 1 h, the bottoms of the filter plates were sealed with Unifilter-96 backing seal tape (PerkinElmer), and 50 µL of MicroScint-O scintillation cocktail (6013611, PerkinElmer) was added to each well. The tops of the plates were then sealed with TopSeal-A sealing film (6050185, PerkinElmer). Radioactivity was measured using a MicroBeta2 reader (PerkinElmer).

Percent inhibition was calculated as % inhibition = [1 − (assay well − Average_LC)/(Average_HC − Average_LC)] × 100. IC_50_ values were determined using the “log(inhibitor) vs. response − variable slope” model in GraphPad Prism 5. Ki values were calculated from IC_50_.

### Spared nerve injury (SNI)-induced mechanical allodynia model in mice

The study was conducted at WuXi AppTec according to its standardized *in vivo* pharmacology protocol for the spared nerve injury (SNI) model. All animal procedures were reviewed and approved by the Institutional Animal Care and Use Committee (IACUC) of WuXi AppTec and were conducted in accordance with applicable institutional guidelines for the care and use of laboratory animals. Male C57BL/6J mice were acclimated for 3 days before surgery. Animals were anesthetized with Zoletil 50 (50 mg/kg) and xylazine (8 mg/kg, i.p.) and subjected to unilateral SNI surgery, in which the tibial and common peroneal nerves were transected while the sural nerve was left intact. Sham-operated mice underwent the same surgical procedure without nerve transection. On day 13 after surgery, animals with a paw withdrawal threshold (PWT) < 0.6 g were randomized into treatment groups (n = 8 per group). Test compounds were administered at a dosing volume of 10 mL/kg. The vehicle consisted of 20% hydroxypropyl-β-cyclodextrin (HP-β-CD). D12-6223 (2 mg/kg), D12-884990 (2 mg/kg), D12-992047 (2 mg/kg), D12-2386724 (2 mg/kg), and D12-1129389-II (1 mg/kg) were administered orally, and PWT was measured at baseline and 1 h after dosing. Analgesic efficacy was expressed as the percentage of maximum possible effect (%MPE), as described below for the mechanical allodynia assessment.

Mechanical allodynia testing was performed at WuXi AppTec according to its standardized *in vivo* pharmacology protocol for pain behavioral assessment. Mechanical sensitivity was assessed using the von Frey up-down method by an experimenter blinded to treatment allocation. Mice were acclimated on a wire-mesh platform for 15 min before testing. Touch-Test Sensory Evaluators (58011, North Coast) were used for stimulation. Calibrated von Frey filaments (0.02, 0.04, 0.07, 0.16, 0.4, 0.6, 1, and 1.4 g for mice) were applied perpendicularly to the lateral plantar surface of the hind paw for 6-8 s at 5-s intervals. A positive response was defined as a sharp paw withdrawal or immediate flinching. If ambulation occurred during stimulation, the stimulus was repeated. Mechanical sensitivity was quantified as the 50% paw withdrawal threshold (PWT), calculated using the formula: 50% response threshold (g) = 10 ^(Xf^ ^+ kδ)^/10,000, where Xf is the logarithmic value of the final von Frey filament used, k is the tabular value corresponding to the pattern of positive and negative responses, and δ is the mean difference, in log units, between consecutive stimuli. Analgesic efficacy was calculated as %MPE = [(Test − Vehicle)/(Sham − Vehicle)] × 100. Statistical significance between pre-dose and post-dose measurements was determined using a paired t-test and figures were generated using OriginPro.

## Author Contributions

Z.C. and L.Z. conceptualized the study. C.M. and Z.C. designed the methodology. C.M. performed the computational enumeration, chemical-space analysis, and PCA. Z.C. and L.Z. designed the biological studies, oversaw their execution. Z.C. and C.M. analyzed the biological data. Z.C. and C.M. wrote the original draft. C.M., Z.C., and L.Z. reviewed and edited the manuscript. Z.C. supervised the study. All authors approved the final version of the manuscript.

## References

1. T. Kamibayashi, M. Maze, Clinical uses of alpha2 -adrenergic agonists. Anesthesiology 93, 1345–1349 (2000).

2. E. A. Fink et al., Structure-based discovery of nonopioid analgesics acting through the α(2A)-adrenergic receptor. Science 377, eabn7065 (2022).

3. J. A. Giovannitti, Jr., S. M. Thoms, J. J. Crawford, Alpha-2 adrenergic receptor agonists: a review of current clinical applications. Anesth Prog 62, 31–39 (2015).

4. J. C. Eisenach et al., Epidural clonidine analgesia for intractable cancer pain. The Epidural Clonidine Study Group. Pain 61, 391–399 (1995).

5. S. Bhatnagar, S. Mishra, S. Madhurima, M. Gurjar, A. S. Mondal, Clonidine as an analgesic adjuvant to continuous paravertebral bupivacaine for post-thoracotomy pain. Anaesth Intensive Care 34, 586–591 (2006).

6. J. E. Hall, T. D. Uhrich, J. A. Barney, S. R. Arain, T. J. Ebert, Sedative, amnestic, and analgesic properties of small-dose dexmedetomidine infusions. Anesth Analg 90, 699–705 (2000).

7. Z. Cheng, Y. Xu, L. Zhu, Targeting peripheral α2A-adrenoceptors as a dual-benefit approach for integrated cancer therapy and supportive care. Cancer Res 86, 5121 (2026).

8. Z. Cheng, C. Moore, H. Li, Y. Xu, L. Zhu, Discovery of a peripherally restricted alpha(2A)AR agonist as a safe and broad-spectrum oral analgesic. Cell Rep Med 7, 102864 (2026).

9. X. Lin et al., A peripheral CB2 cannabinoid receptor mechanism suppresses chemotherapy-induced peripheral neuropathy: evidence from a CB2 reporter mouse. Pain 163, 834–851 (2022).

10. V. Tiwari et al., Peripherally Acting mu-Opioid Receptor Agonists Attenuate Ongoing Pain-associated Behavior and Spontaneous Neuronal Activity after Nerve Injury in Rats. Anesthesiology 128, 1220–1236 (2018).

11. V. A. Rangari et al., A cryptic pocket in CB1 drives peripheral and functional selectivity. Nature 640, 265–273 (2025).

12. N. Vadivelu, S. Mitra, R. L. Hines, Peripheral opioid receptor agonists for analgesia: a comprehensive review. J Opioid Manag 7, 55–68 (2011).

13. L. Zhu, Z. Cheng, W. I. P. O. (WIPO), Ed. (AlpheraBio LLC, WO, 2024), vol. A2, chap. WO2024226608A2.

14. L. Zhu, Z. Cheng, W. I. P. O. (WIPO), Ed. (AlpheraBio LLC, WO, 2025), vol. A1, chap. WO2025171183A1.

15. H. Zhao, Z. Guo, Medicinal chemistry strategies in follow-on drug discovery. Drug Discov Today 14, 516–522 (2009).

16. J. Eder, R. Sedrani, C. Wiesmann, The discovery of first-in-class drugs: origins and evolution. Nat Rev Drug Discov 13, 577–587 (2014).

17. L. Peltason, N. Weskamp, A. Teckentrup, J. Bajorath, Exploration of structure-activity relationship determinants in analogue series. J Med Chem 52, 3212–3224 (2009).

18. D. M. Parry, Closing the Loop: Developing an Integrated Design, Make, and Test Platform for Discovery. ACS Med Chem Lett 10, 848–856 (2019).

19. There’s room for a little randomness in drug development. Medchemcomm 8, 1738 (2017).

20. C. Lipinski, A. Hopkins, Navigating chemical space for biology and medicine. Nature 432, 855–861 (2004).

21. J. L. Reymond, M. Awale, Exploring chemical space for drug discovery using the chemical universe database. ACS Chem Neurosci 3, 649–657 (2012).

22. C. J. Gerry, S. L. Schreiber, Chemical probes and drug leads from advances in synthetic planning and methodology. Nat Rev Drug Discov 17, 333–352 (2018).

23. G. Schneider, Virtual screening: an endless staircase? Nat Rev Drug Discov 9, 273–276 (2010).

24. T. Yang et al., DrugSpaceX: a large screenable and synthetically tractable database extending drug space. Nucleic Acids Res 49, D1170–D1178 (2021).

25. D. Montes-Grajales, L. Menestrina, R. Garcia-Serna, J. Mestres, ChemBang: Expanding the Chemical Space Around Small Molecules. Mol Inform 45, e70036 (2026).

26. M. Segall et al., Applying medicinal chemistry transformations and multiparameter optimization to guide the search for high-quality leads and candidates. J Chem Inf Model 51, 2967–2976 (2011).

27. https://optibrium.com/products/stardrop

28. D. Rogers, M. Hahn, Extended-connectivity fingerprints. J Chem Inf Model 50, 742–754 (2010).

29. C. R. Harris et al., Array programming with NumPy. Nature 585, 357–362 (2020).

30. I. T. Jolliffe, J. Cadima, Principal component analysis: a review and recent developments. Philos Trans A Math Phys Eng Sci 374, 20150202 (2016).

31. E. Martin, E. Cao, Euclidean chemical spaces from molecular fingerprints: Hamming distance and Hempel’s ravens. J Comput Aided Mol Des 29, 387–395 (2015).

